# Parent-offspring brain similarity: Specificities and commonalities across gender combinations - the Transmit Radiant Individuality to Offspring (TRIO) study

**DOI:** 10.1101/2024.10.05.616578

**Authors:** Izumi Matsudaira, Ryo Yamaguchi, Yasuyuki Taki

## Abstract

Research suggests that parent-offspring brain similarities may underlie intergenerational transmission of psychopathology. However, most studies have focused on mothers and offspring, with few including fathers. This study aimed to extend understanding of parent-offspring neural similarities by examining parent-offspring trios. The study included 152 Japanese biological parent-offspring trios who participated in the Transmit Radiant Individuality to Offspring (TRIO) study. We analyzed the parent-offspring similarities in brain structural features (cortical thickness, surface area, local gyrification index, and subcortical volume) across different parent-offspring gender combinations (father-son, father-daughter, mother-son, and mother-daughter). Additionally, we investigated the relationship between brain similarities and similarities in intelligence and personality traits in parents and offspring. Our findings confirmed that correlations in brain structural features between father-offspring or mother-offspring dyads were significantly stronger than those between unrelated individuals. Notably, both sons and daughters exhibited brain regions similar to their fathers only, mothers only, both parents, or neither parent. Furthermore, a significant association was observed between similarities in general intelligence and the surface area of auditory regions in both father-offspring and mother-offspring dyads. These results provide valuable insights into the genetic and environmental factors influencing brain development and aging across generations. This study is expected to contribute to future research elucidating the mechanisms underlying the intergenerational transmission of psychiatric disorders.

## Introduction

Intergenerational transmission refers to offspring acquiring behaviors and traits that are similar to those of their parents (1). Several studies have demonstrated the intergenerational continuities of psychopathologies (2–5). Recent large-sample studies indicate that children with a history of parental psychiatric hospitalization are 2-3 times more likely to develop mental illness in adolescence than those without such a history (6). Children exposed to a parental suicide attempt in early childhood are at an increased risk of attempting suicide during adolescence (7). Therefore, the intergenerational transmission of psychopathology is likely one of the key issues in contemporary psychiatry (8). Although intergenerational transmission is believed to occur through gene-environment interactions, the specific mechanisms underlying this process remain poorly understood.

Intergenerational neuroimaging can help explore the mechanism of intergenerational transmission (9). Several groundbreaking studies have employed magnetic resonance imaging (MRI) to investigate the structural and functional brain similarities among parent-offspring pairs. Parent-offspring dyads have been shown to exhibit greater similarity in brain features than random unrelated adult-child pairs (10–13). Parent-offspring similarities have been reported in various brain features, such as in gray matter volume (GMV) (10, 12, 14, 15), cortical thickness (CT) (16), surface area (SA) (11, 15), local gyrification index (LGI) (11), sulcal morphology (11, 13), white matter microstructure (17–19), neural metabolism (20), resting-state functional connectivity (10, 21, 22), and neural activation during reward processing (23). Some of these studies suggest that neural similarities may underlie the intergenerational transmission of mental illness (14–16). However, there is a lack of consensus owing to considerable heterogeneity among these studies with respect to the study population and methodologies (24).

Despite progress in intergenerational neuroimaging, several challenges persist. First, most studies have not examined differences in the parent-offspring similarity across various brain measures, despite their distinct developmental trajectories. Research on human brain development has primarily focused on CT, SA, subcortical volume (SV), and indices of cortical folding (gyrification). Each of these characteristics matures at different times (25–30) and has a distinct genetic background (31–33). Comparing parent-offspring similarities for each characteristic within the same sample can provide novel insights. Second, the relationship between neural similarities and the intergenerational transmission of specific behavior remains unclear. To date, only one study has explored the association between mother-offspring neural and behavioral similarities, revealing that similarities in well-being are predicted by similarities in GMV in the anterior cingulate cortex (ACC) and prefrontal cortex (12). Third, most previous studies have focused on brain regions involved in higher-order functions (11, 12, 14, 15), leaving a significant knowledge gap regarding parent-offspring similarities in other brain regions (24). Fourth, many of the offspring studied in previous research were in early childhood to adolescence (10–14, 16, 17, 19, 22, 23), leaving a dearth of knowledge on parent-child neural similarity at ages when brain maturation has progressed. Fifth, most importantly, contemporary research has largely focused on mother-offspring dyads (11–13, 16, 22, 23), while data on fathers, even when available, are often very limited (10, 17–19, 21). Genetic studies have revealed that genomic imprinting, an epigenetic phenomenon in which certain genes are expressed from either the maternally or paternally inherited allele (34), occurs more frequently in the brain than in somatic tissues (35, 36). These parent-of-origin effects on brain development vary depending on offspring’s sex (37). Hence, it is essential to investigate the neural similarities between both mother-offspring and father-offspring dyads. Although pioneering studies have investigated neural similarities across mother-daughter, mother-son, father-daughter, and father-son dyads (14, 15), these studies lacked comparisons with unrelated adult-child pairs, leaving the assumption that similarities are specific to parent-offspring dyads unverified (24). Therefore, a new study using balanced datasets of father-offspring and mother-offspring pairs is needed to overcome the aforementioned limitations.

This study aimed to elucidate the specificities and commonalities in parent-offspring similarities of brain structural characteristics (CT, SA, LGI, and SV) among different parent-offspring gender combinations, using parent-offspring trios consisting of offspring from adolescence to young adulthood and their biological parents (38). Additionally, we examined gender-specific differences in the behavioral similarities between parents and offspring, focusing on intelligence and personality traits that can be assessed using the same scale across a broad age range. To investigate whether neural similarity underlies the intergenerational transmission of behavior, we also tested the association between neural and behavioral similarities. We hypothesized that mother-specific or father-specific neural and behavioral similarities are observed in both sons and daughters.

## Materials and methods

### Participants

The study population comprised 152 parent-offspring trios consisting of three members: male and female offspring aged 15 to 40 and their biological father and mother aged 65 and younger. This research was part of an ongoing project investigating the intergenerational transmission effect on brain development (38). To control for genetic background, participation was limited to Japanese individuals with no relatives within the third degree of kinship from other ethnicities. Individuals with a history of cerebrovascular disease, brain tumor, intracranial disease, degenerative brain disease, epilepsy, severe heart disease, and brain injury with impaired consciousness were not eligible for inclusion. These conditions were verified at the time of participation, and if any trio member met the exclusion criteria, their participation was declined. This study adhered to the principles enshrined in the Declaration of Helsinki (39) and received approval from the Institutional Review Board of Tohoku University (Approval No. 2022-1-534). Written informed consent was obtained from all participants before the study. For minors (age <18 years), parental consent was also required.

Following rigorous screening, two parent-offspring trios were excluded due to the offspring’s neurological disorders and suboptimal brain imaging quality. Additionally, five fathers were excluded from the study for the following reasons: two exceeded the upper age limit for inclusion, one had a neurological disorder, one had a brain hematoma, and one declined to undergo MRI scanning. Finally, 145 father-offspring dyads (58 sons and 87 daughters) and 150 mother-offspring dyads (60 sons and 90 daughters) were included in this study.

### Neuroimaging

Brain images were acquired using a 3-Tesla dStream Achieva scanner (Philips Medical Systems, Best, Netherlands) equipped with a 20-channel head-neck coil. For each participant, two types of images were acquired. Sagittal T1-weighted images (T1WIs) were acquired using a magnetization-prepared rapid gradient-echo sequence with parameters including 368 × 368 matrix, 11 ms repetition time, 5.1 ms echo time, 256 × 256 mm field of view, 257 slices, and 0.7 mm slice thickness. Sagittal T2-weighted images (T2WIs) were acquired using a spin-echo sequence with parameters including 368 × 368 matrix, 2500 ms repetition time, 3200 ms echo time, 256 × 256 mm field of view, 250 slices, and 0.7 mm slice thickness. Image quality was visually inspected immediately after imaging, and if significant motion artifacts were detected, re-imaging was performed.

### Preprocessing of brain images

All neuroimaging data was preprocessed using FreeSurfer v7.3.2 to estimate CT, SA, and LGI of cortical regions, and SV (40–42). To enhance segmentation quality, three steps preceded the standard recon-all pipeline. First, AC-PC alignment was conducted using a 6-degree-of-freedom rigid body transformation (43). Second, the nonuniform intensity normalization was conducted using N4ITK (44). Third, brain extraction was performed using HD-BET (45). These three processes were applied to both T1WIs and T2WIs. Additionally, the brain mask image generated by HD-BET for T1WIs was utilized by incorporating the ‘-xmask’ option in the “recon-all” process. As HD-BET removed extracerebral tissue, the ‘-noskullstrip’ option was used to prevent skull stripping within the recon-all pipeline. Quality assurance of the preprocessing results was conducted through visual inspection and manual editing of the reconstructed images. Specifically, we verified that the boundaries between white and gray matter, as well as the pial surface, were accurately delineated. Any images with segmentation errors or topological defects underwent manual corrections, followed by a re-run of the recon-all process. CT, SA, and SV were obtained from the standard recon-all output. To obtain LGI, we performed an additional recon-all run with the ‘-localGI’ flag (42). This process was performed additionally after the completion of quality assurance.

CT, SA, and LGI for each brain region were calculated based on the Human Connectome Project Multi-Modal Parcellation version 1.0 (HCP-MMP1) atlas (46). However, since the HCP-derived atlas and the FreeSurfer data are on different surface spaces (fs_LR mesh and fsaverage mesh, respectively), we resampled the HCP-MMP1 label map file downloaded from the BALSA database (https://balsa.wustl.edu/file/L632n) (47) into individual surface data, using Connectome Workbench version 1.5.0(http://www.humanconnectome.org/software/connectome-workbench.html). The resampling procedure was based on the document in the HCP Public Pages (https://wiki.humanconnectome.org/docs/assets/Resampling-FreeSurfer-HCP_5_8.pdf).Additionally, SV was calculated based on the Automatic Subcortical Segmentation (aseg) atlas (41).

### Behavioral assessments

Wechsler Adult Intelligence Scale Fourth Edition (WAIS-IV) (48) was used to measure the participants’ global intelligence. For offspring aged 15, the Wechsler Intelligence Scale for Children Fourth Edition (WISC-IV) (49) was used. Ten core subtests were conducted to calculate the full-scale intelligence quotient (FSIQ) along with four factors (verbal comprehension index [VCI]; perceptual reasoning index [PRI]; working memory index [WMI]; and processing speed index [PSI]). These five scores were used for the subsequent analysis. The tests were conducted in a quiet, distraction-free room, one-on-one with trained staff familiar with the procedure and participants.

Personality traits were assessed using the NEO five-factor inventory (NEO-FFI) (50), a 60-item questionnaire that evaluates five personality traits (neuroticism, extraversion, openness to experience, agreeableness, conscientiousness) using a five-point Likert scale. A previous study has demonstrated the reliability and validity of the Japanese version of the NEO-FFI (51).

### Statistical analysis

#### Similarities in CT, SA, LGI, and SV

The analysis of parent-offspring brain similarity was conducted following the procedures suggested in previous studies (11, 12). Here, we explain the process using CT similarity between mother and offspring as an example. Pearson’s correlation coefficient was used as the index of similarity. To determine if the CT in a specific brain region was more similar between mothers and offspring than between unrelated individuals, we employed a permutation approach using MATLAB (MathWorks Inc., Natick, MA, USA). This involved randomly re-pairing mothers with offspring to create unrelated pairs, and then calculating the Pearson’s correlation coefficient for CT of that region. This process was repeated 1,000 times, generating a normal distribution of correlation coefficients for unrelated individuals. In the analysis of CT or LGI, the offspring’s age was controlled for. For SA and SV analyses, both the offspring’s age and total intracranial volume (TIV) of the parent and offspring were controlled for. All 1,000 correlation coefficients were Fisher’s z-transformed, averaged, and then back-transformed into a correlation coefficient, yielding a similarity score for unrelated pairs. This score was compared to the correlation coefficient of real mother-offspring dyads. If the correlation coefficient of real dyads was significantly higher, the CT of that region was deemed similar between mother and offspring. The “cocor” package in R was used to test the differences in correlation coefficients (52). For CT, SA, and LGI, similarity was examined in 360 regions based on the HCP-MMP1 atlas. For SV, similarity was assessed in 32 regions, including the hippocampus, amygdala, thalamus, basal ganglia, cerebellum, brainstem, ventricles, and corpus callosum. To account for multiple comparisons, the False Discovery Rate (FDR) was adjusted using the Benjamini-Hochberg (BH) method. A statistical threshold of FDR-corrected *p* (*q* value) < 0.05 was established to determine significant brain similarity between parents and offspring.

After conducting the above analyses separately for father-offspring and mother-offspring pairs, we also conducted them for each parent-offspring gender combination (father-daughter, father-son, mother-daughter, mother-son).

### Similarities in intelligence and personality traits

The similarity between parents and offspring in intelligence and personality traits was examined using the same procedure employed for brain structure similarity. The offspring’s age was controlled for in the correlation analysis. Parent-offspring dyads with missing scores for either the parent or offspring were excluded. The final sample size for the analysis of intelligence was 140 father-offspring dyads (56 sons, 84 daughters) and 147 mother-offspring dyads (59 sons, 88 daughters). For personality traits, the sample sizes were 142 father-offspring dyads (56 sons, 86 daughters) and 148 mother-offspring dyads (58 sons, 90 daughters). To account for multiple comparisons, the FDR was adjusted using the BH method, with a statistical threshold of *q* < 0.05. Analyses were conducted separately for father-offspring dyads, mother-offspring dyads, and each parent-offspring gender combination, mirroring the approach used in the neuroimaging analysis.

### The association between brain structure similarity and behavioral similarity

The Euclidean distance between parent and offspring was used to measure the extent of similarity in brain regions and behavioral indices (intelligence and personality traits) where similarity was observed in the previous analysis. In both cases, the Euclidean distance was calculated using standardized values of the raw measurements. Multiple regression analysis was conducted using the Euclidean distance of behavioral indicators as a dependent variable, the Euclidean distance of brain features as the predictor, and the offspring’s age as a covariate. When SA and SV were used as predictors, the TIV of both the parent and offspring was added to the model as a covariate. For each of CT, SA, LGI, and SV, the FDR was adjusted using the BH method, with a statistical threshold set at *q* < 0.05.

## Results

### Demographic information

Descriptive statistics for participants’ age, intelligence, and personality traits are presented in Table 1.

**Table 1.**
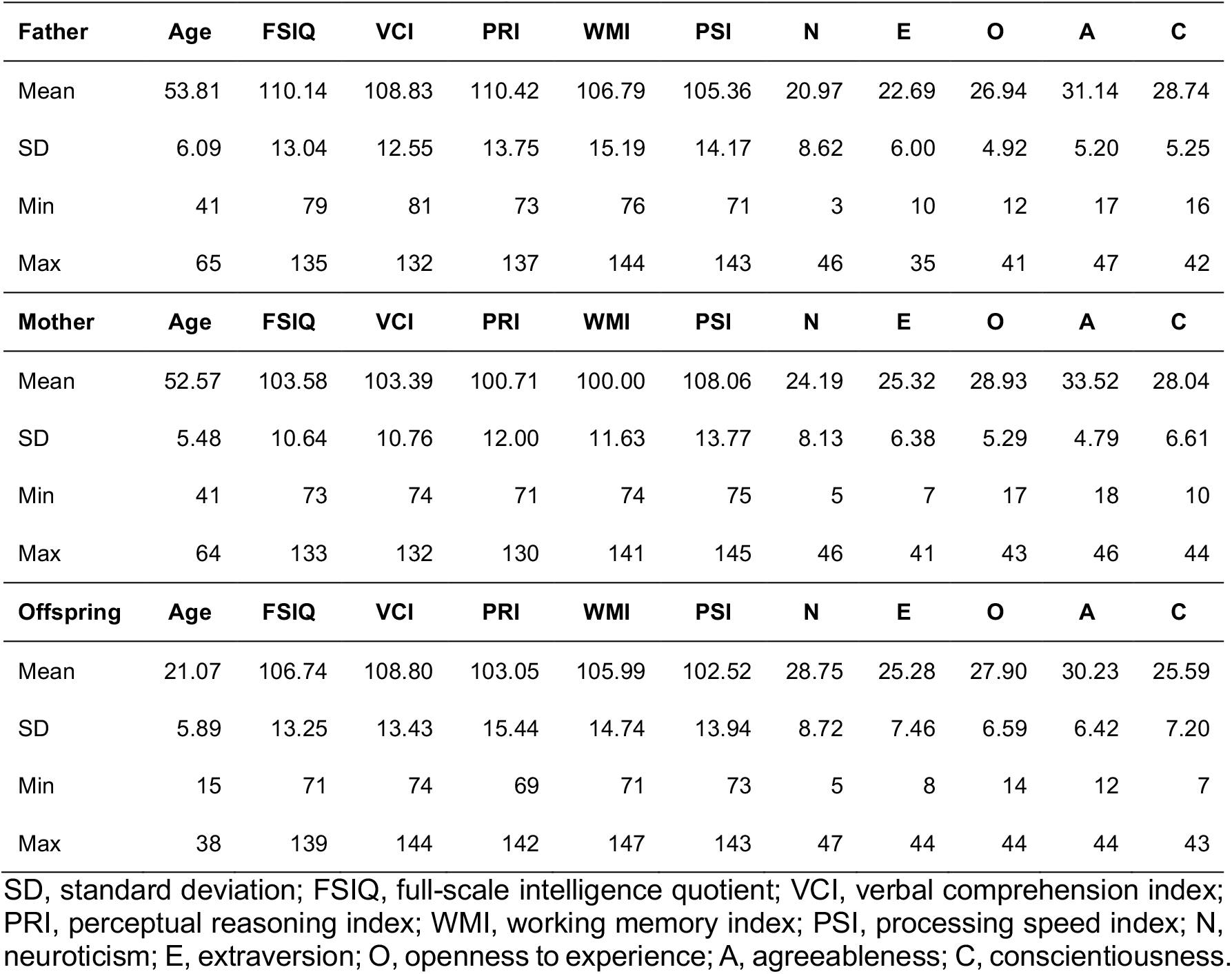
Characteristics of the study population.

### Neural similarities in father-offspring dyads and mother-offspring dyads

Neural similarities between father-offspring dyads and/or mother-offspring dyads were observed across various brain regions. Figure 1 illustrates the brain regions where neural similarity exceeded chance levels (*q* < 0.05), indicating a significant difference in correlation coefficients between parent-offspring dyads and unrelated pairs. The detailed results for all brain regions are provided in Supplementary Tables 1-8. Notably, CT analysis revealed that 150 out of 360 regions in father-offspring pairs and 149 regions in mother-offspring pairs showed significantly greater similarity compared to unrelated pairs (Supplementary Tables 1 and 5). A particularly strong similarity was observed in the right 3a (primary somatosensory cortex) for father-offspring (Pearson’s *r* in real dyads = 0.593, averaged Pearson’s *r* in unrelated pairs = -0.002, *z* = 5.767, *q* < 0.001) and mother-offspring dyads (Pearson’s *r* in real dyads = 0.539, averaged Pearson’s *r* in unrelated pairs = -0.002, *z* = 5.174, *q* < 0.001). In addition to the right primary somatosensory cortex, the left primary somatosensory cortex also exhibited strong similarity in mother-offspring pairs (Pearson’s *r* in real dyads = 0.554, averaged Pearson’s *r* in unrelated pairs = -0.002, *z* = 5.382, *q* < 0.001). For SA, 65 out of 360 regions in father-offspring pairs and 156 regions in mother-offspring pairs showed significantly greater similarity compared to unrelated pairs (Supplementary Tables 2 and 6). Notably, bilateral visual cortex, insular cortex, and frontal operculum regions exhibited similar neural features in both father-offspring and mother-offspring dyads. Mother-offspring dyads, however, displayed particularly strong similarities in the somatosensory cortex and cingulate cortex. LGI analyses revealed widespread similarities between both father-offspring and mother-offspring dyads in almost all regions (Supplementary Tables 3 and 7). The sole exception was the left 7AL (lateral area 7A) in father-offspring dyads (Pearson’s *r* in real dyads = 0.199, averaged Pearson’s *r* in unrelated pairs = -0.001, *z* = 1.712, *q* = 0.086), which did not show a significant difference in correlation coefficients compared to unrelated pairs. In SV analyses, the similarity in the volume of bilateral cerebellar white matter was common between father-offspring and mother-offspring dyads (Supplementary Tables 4 and 8).

**Figure 1.**
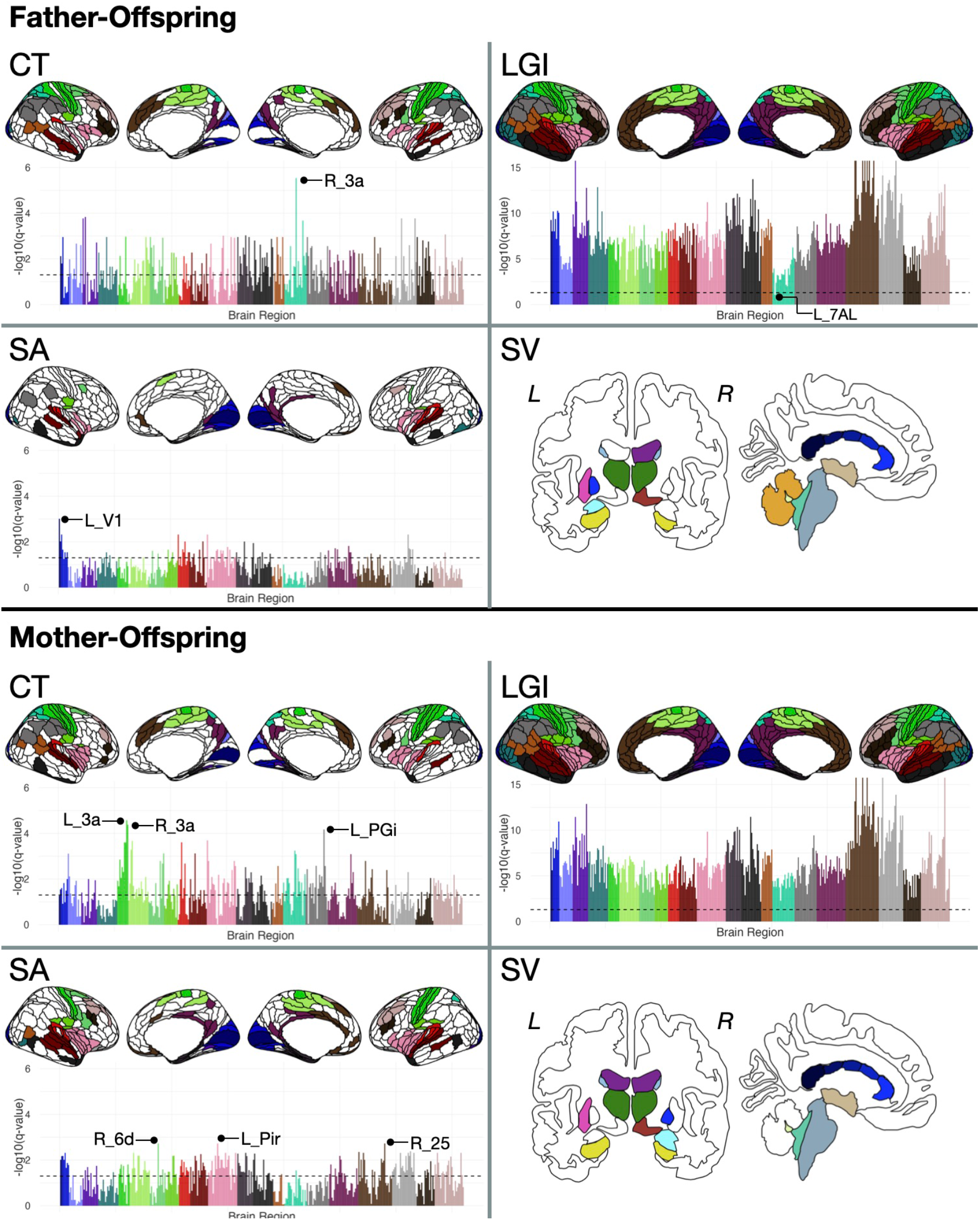
Neural similarities in father-offspring dyads and mother-offspring dyads. The original HCP-MMP1 atlas (46) comprises 180 regions per hemisphere, grouped into 22 larger sections. The regions for which significantly stronger correlations were observed in father-offspring or mother-offspring dyads compared to unrelated pairs are color-coded. The *q*-values for the 360 brain regions are shown in the plot. The vertical axis of the plot represents the log-transformed *q*-values. Each bar represents an individual brain region, and the colors of the bars correspond to those in the brain illustration. Subcortical regions where significantly stronger correlations between parents and offspring compared to unrelated pairs were observed are color-coded according to the ggseg plotting tool (53). CT, cortical thickness; SA, surface area; LGI, local gyrification index; SV, subcortical volume, L, left hemisphere; R, right hemisphere.

### Differences in neural similarities based on the parent-offspring gender combination

Based on the analysis of father-son, father-daughter, mother-son, and mother-daughter dyads (Supplementary Figures 1 and 2, Supplementary Tables 9-24), regions were colored to indicate those where sons and daughters showed similarity exclusively with the mother, exclusively with the father, with both parents or with neither parent (Fig.2). For CT, notable gender-specific patterns emerged. Daughters exhibited unique similarities with their fathers in several regions (Supplementary Table 17). Sons, however, showed similarity with both parents in the right 3a (father-son; Pearson’s *r* in real dyads = 0.659, averaged Pearson’s *r* in unrelated pairs = -0.002, *z* = 4.154, *q* = 0.012, mother-son; Pearson’s *r* in real dyads = 0.658, averaged Pearson’s *r* in unrelated pairs = -0.016, *z* = 4.319, *q* = 0.003) but did not exhibit exclusive similarity with their fathers in any region. For SA, daughters showed similarity only with their mothers in regions corresponding to the superior temporal gyrus (STG) and orbitofrontal cortex (OFC) (Supplementary Table 22). The primary visual cortex showed similarity with the mother only on the right side (Pearson’s *r* in real dyads = 0.449, averaged Pearson’s *r* in unrelated pairs = 0.001, *z* = 3.177, *q* = 0.047) and with the father only on the left side (Pearson’s *r* in real dyads = 0.527, averaged Pearson’s *r* in unrelated pairs = -0.005, *z* = 3.836, *q* = 0.017). Sons did not show similarity with either parent in any of the regions (Supplementary Tables 10 and 14). For LGI, numerous brain regions exhibited similarity with both parents, regardless of the offspring’s gender. However, several regions showed gender-specific similarities. In daughters, the left LIPv (lateral inferior parietal complex), 7AL, and VIP (ventral intraparietal complex) showed exclusive similarity to mothers (LIPv; Pearson’s *r* in real dyads = 0.405, averaged Pearson’s *r* in unrelated pairs = -0.007, *z* = 2.882, *q* = 0.004, 7AL; Pearson’s *r* in real dyads = 0.449, averaged Pearson’s *r* in unrelated pairs = -0.001, *z* = 3.194, *q* = 0.002, VIP; Pearson’s *r* in real dyads = 0.362, averaged Pearson’s *r* in unrelated pairs = -0.005, *z* = 2.536, *q* = 0.011), while the left 47l (Pearson’s *r* in real dyads = 0.496, averaged Pearson’s *r* in unrelated pairs = -0.003, *z* = 3.547, *q* < 0.001) was similar only to the father. In sons, the somatosensory cortex, inferior frontal gyrus, superior temporal sulcus, and auditory cortex showed exclusive similarity with fathers, while part of the lateral temporal cortex was similar only to mothers (Supplementary Table 15). For SV, bilateral hippocampus and caudate showed a gender-specific association, with sons exhibiting similarity exclusively with their fathers, and daughters exhibiting similarity exclusively with their mothers (Supplementary Tables 12 and 24). In contrast, the right lateral ventricle volume displayed a cross-gender parental influence, with sons resembling their mothers and daughters resembling their fathers (Supplementary Tables 16 and 20).

**Figure 2.**
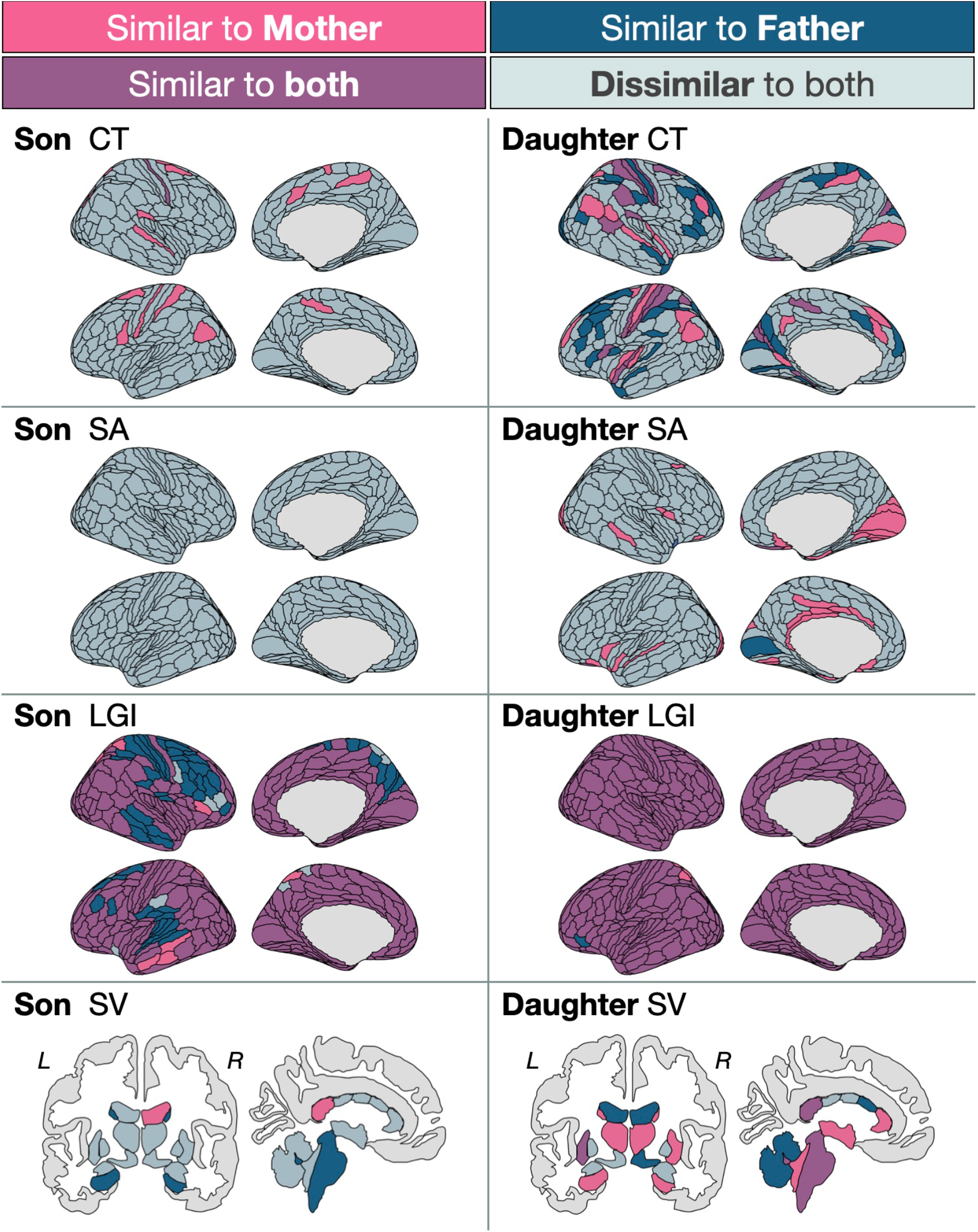
Comparison of neural similarities among parent-offspring gender combinations. Each brain region is color-coded based on the comparison of correlation coefficients with unrelated pairs: pink if significant only in mother-offspring pairs, navy blue if significant only in father-offspring pairs, purple if significant in both mother-offspring and father-offspring pairs, and gray if nonsignificant in both. CT, cortical thickness; SA, surface area; LGI, local gyrification index; SV, subcortical volume, L, left hemisphere; R, right hemisphere.

### Association between neural similarities and behavioral similarities

Figure 3A illustrates the intelligence and personality traits that exhibited significant parental-offspring correlations, exceeding those of unrelated pairs (see Supplementary Figures 3-5, Supplementary Tables 25-36 for detailed results). Similarities in FSIQ and VCI were observed between fathers and their offspring, with this pattern holding specifically for sons. Similarities in FSIQ, VCI, WMI, and conscientiousness were observed between mother-offspring dyads. Similarities in FSIQ, VCI, and WMI were observed between mothers and sons, while the similarity in conscientiousness was observed between mothers and daughters.

**Figure 3.**
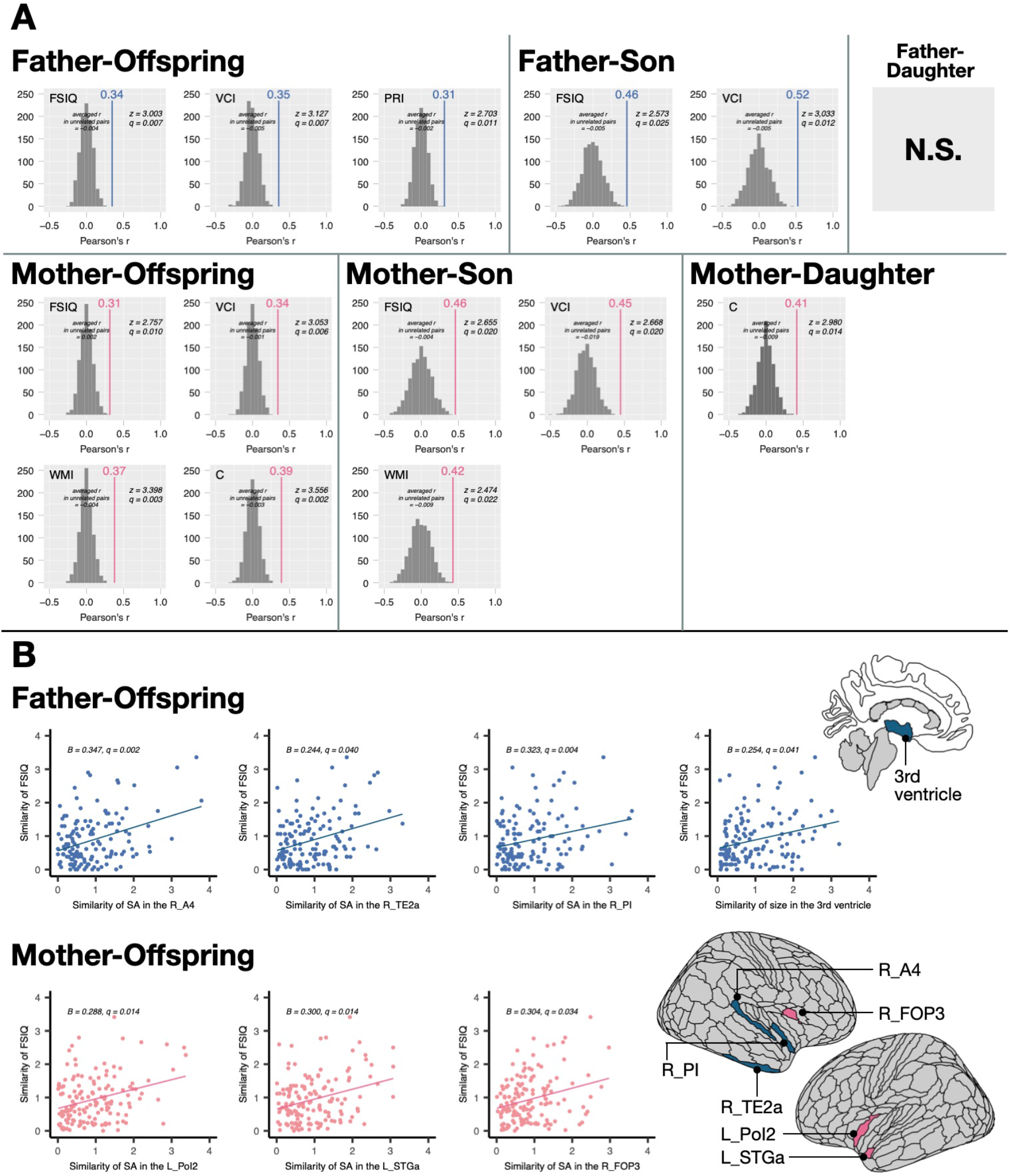
Parent-offspring behavioral similarities. (A) The differences in correlation coefficients between real parent-offspring pairs and unrelated pairs are shown for father-offspring, mother-offspring, and each gender combination. The gray histogram represents the distribution of correlation coefficients for 1,000 patterns of unrelated pairs. “averaged r in unrelated pairs” means Z-transformed, averaged, and backtransformed correlation coefficients of unrelated pairs. The correlation coefficients for real parent-offspring pairs are shown as solid lines, with the corresponding values labeled on top. FSIQ, full-scale intelligence quotient; VCI, verbal comprehension index; WMI, working memory index; C, conscientiousness. (B) Scatter plots show the relationship between the Euclidean distance for FSIQ and the Euclidean distance for each brain region in father-offspring and mother-offspring dyads. The standardized partial regression coefficients and *q*-values are displayed in the top left corner of the scatter plot. SA, surface area; L, left hemisphere; R, right hemisphere.

Figure 3B illustrates the relationships between neural similarities and behavioral similarities (Fig.3B). Notably, the similarity in FSIQ between father-offspring dyads was significantly and positively predicted by similarities in the volume of the third ventricle, as well as the SA in the right A4 (auditory 4 complex), TE2a (temporal area 2 anterior), and PI (para-insular area). The similarity in FSIQ between mother-offspring dyads was significantly predicted by similarities in the SA of the left PoI2 (posterior insular area 2), STGa (superior temporal gyrus anterior), and FOP3 (frontal opercular area 3). However, when analyzing specific parent-offspring gender combinations, no significant associations were observed between neural and behavioral similarities.

## Discussion

This study reveals that parent-offspring brain similarities vary across different gender combinations. Our findings also indicate that similarities in FSIQ between parents and offspring are correlated with similarities in the SA of specific brain regions, underscoring the potential neuroanatomical basis of cognitive inheritance.

Our findings support the notion that daughters’ brains resemble their parents more than sons’, consistent with a previous study (10). Daughters exhibited similarities with their fathers and/or mothers in numerous brain regions, compared to sons. This suggests a robust tendency for daughters to dominate neural similarities with parents. Previous studies have suggested matrilineal transmission patterns in specific brain structures such as GMV in the ACC, ventromedial prefrontal cortex (vmPFC), and caudal middle frontal gyrus (MFG) (14, 15), and SA in the caudal MFG (15). In our sample, SA in the ACC (a24pr, anterior 24 prime; p24pr, posterior 24 prime) was similar only between mother-daughter dyads. Cortical GMV is a measure of the combination of CT and SA, and the statistical results of GMV are mainly influenced by SA (54). Thus, our results are partially consistent with previous studies. However, SA in the right OFC, which is identical to vmPFC, was also similar in father-daughter dyads, not only in mother-daughter dyads. Utilizing trio samples allowed us to identify these novel similarities in regions previously considered mother-daughter specific, expanding our understanding of parent-offspring brain similarities and highlighting the importance of considering both maternal and paternal influences.

Our analysis reveals evidence of a potential parent-of-origin effect in the development of certain brain regions. For both sons and daughters, the CT of the left PGi (PG inferior), right 5mv (area 5m ventral), and right A5 (auditory 5 complex) exhibited similarity only with their mothers, while the LGI of the left 7AL and left VIP exhibited similarity only to their fathers. The volumes of the bilateral hippocampus and caudate nucleus showed similarities only between daughters and their mothers, and between sons and their fathers, suggesting that the developmental processes in these regions may share specific commonalities between same-sex parent-offspring dyads. On the other hand, specific brain regions exhibited similarities between opposite-sex parent-offspring dyads. The right lateral ventricle volume and the CT of the left 6a (area 6 anterior) and right IP0 (area intraparietal 0) were similar only between daughters and their fathers and between sons and their mothers. These findings suggest unique commonalities in brain development between opposite-sex parent-offspring dyads. Previous genetic studies have elucidated the influence of genomic imprinting on brain development (55). In the mouse brain, more than 1,300 loci with maternal or paternal allelic expression bias have been detected (56). Specifically, gene expression from the maternally inherited X chromosome is preferentially observed in the cortex of female offspring, while that from the paternally inherited X chromosome is preferentially observed in the hypothalamus (37). Although it remains unclear whether and how these phenomena are related to human brain development, it is possible that genomic imprinting contributes to the variation in parent-offspring brain similarities depending on sex combinations. Additionally, since both CT and SV exhibit experience-dependent plasticity (57–59), it is also possible that the emotional and physiological responses shared between parents and offspring contribute to these similarities. Previous research has demonstrated neural synchrony between parents and offspring during tasks or rest, although offspring in those studies were younger than those in our sample (60–63). Further study is required to examine the relationship between brain structural similarities, genetic factors, and environmental factors based on parent-offspring gender combinations.

Our findings revealed an unexpected complexity in parent-offspring brain similarities, with certain regions showing resemblance to both parents and others showing no similarity to either. Notably, the CT of the right 3a exhibited similarity between both sons and daughters and their parents. These findings suggest that the development of this region may be influenced by the relationship among the father, mother, and offspring, regardless of gender.

Surprisingly, the LGI of most brain regions in both sons and daughters was similar to that of their parents. This finding suggests that the LGI of couples may also be similar. Gyrification is an indicator of cortical folding that is largely determined during fetal development and remains relatively stable after birth (64, 65). Therefore, our results suggest that assortative mating at the level of gyrification may be involved in the brain development of offspring. Additionally, there were many regions where CT and/or SA were not similar to either parent, especially in sons. This observation suggests that factors beyond parental influence may shape brain development. Interestingly, research has shown that children’s brain structures can resemble those of close friends (66), implying a potential role for social interactions outside of the parent-child relationship.

Intelligence and personality traits showed similarities between parents and offspring, but these behavioral similarities did not necessarily correlate with brain structural similarities. However, similarities in FSIQ between father-offspring and mother-offspring dyads were linked to similarities in SA of regions involved in auditory and language functions. The specific regions associated with FSIQ similarity differed between dyads. When analyzing offspring by gender, this association was not observed, which may be attributed to relatively small sample sizes (60 sons and 90 daughters), rather than indicating a genuinely gender-independent relationship. Behavioral measures, such as FSIQ, involve the activation of many cognitive functions, making it unlikely that structural similarities in individual brain regions can fully account for behavioral similarities. A more detailed understanding may be achieved by examining the relationship between parent-offspring similarities in functional or structural connectivity and behavioral similarities. Also, since these regions are susceptible to distortion due to the inhomogeneity of MR signals, these findings need to be replicated.

Some limitations of this study should be considered while interpreting the findings. First, the differing sample sizes between sons and daughters may impact the detection of gender-specific effects, potentially masking subtle differences due to effect size. Additionally, the greater propensity of daughters to participate in activities with their parents highlights a recruitment bias. To address this, methodological innovations are needed to boost the participation rates of sons and ensure more representative cohorts in future research. Second, the wide age range among offspring may introduce age-related brain changes that cannot be fully controlled for, even with age as a covariate, in statistical analysis. As most offspring in this study were teenagers, future research should recruit more participants in their 20s and 30s and conduct age-group-specific analyses. Third, this study’s cross-sectional design limits the understanding of development trajectories. Long-term longitudinal studies are required to clarify when and how parent-offspring similarities in brain structure and behavior emerge and evolve.

In conclusion, our study using Japanese trio samples confirmed that brain structure and behavior similarities exist between parents and offspring, exceeding those found between unrelated individuals. Furthermore, we discovered that whether specific brain regions and features resemble the father, the mother, both, or neither varies depending on the offspring’s gender. This study provides valuable insights into brain development and aging in parents and offspring. Our findings are expected to contribute to future investigations into the mechanisms underlying the intergenerational transmission of psychiatric disorders.

## Acknowledgements

We wish to express our deepest thanks to the participants for their willingness to participate in this study. Special thanks go to Yukiko Suginome, Saeko Hoshi, Chieko Miura, Maiko Chiba, Misaki Abe, Junko Kato, and Shuzo Yamamoto for their invaluable support. We are also grateful to Dr. Ryosuke Kimura, Dr. Tadashi Imanishi, Dr. Hiroaki Tomita, Ayaka Uchiyama, Kanna Oyama, Mihiro Koizumi, Takumi Uchiyama, Yuka Aoki, Fumiaki Nitta, Nanae Moriya, Fumi Seki, Narumi Fujiwara, Kazuki Ozawa, Yuji Yanagiya, Mako Toyota, Sawako Watanabe, Yuka Hatayama, Megumi Kato, Maiko Suenaga, Megumi Shirahama, Rinka Yoshihara, Jun Nomura, Ryuhei Ohgi, Ruri Takahashi, Yuto Tanaka, Saki Uchida, and Ruriko Igarashi for their contributions to data collection. Additionally, we extend our gratitude to Dr. Kiyotaka Nemoto and Dr. Takuya Hayashi for their support with the neuroimaging analysis, and to Dr. Plamina Dimanova for sharing the MATLAB script for the permutation analysis. We appreciate all the institutions that cooperated with us in recruiting study participants. Lastly, we would like to thank Enago (https://www.enago.jp/) for English language editing.

