## Supplementary Figures for "Parent-offspring brain similarity: Specificities and commonalities across gender combinations - the Transmit Radiant Individuality to Offspring (TRIO) study"

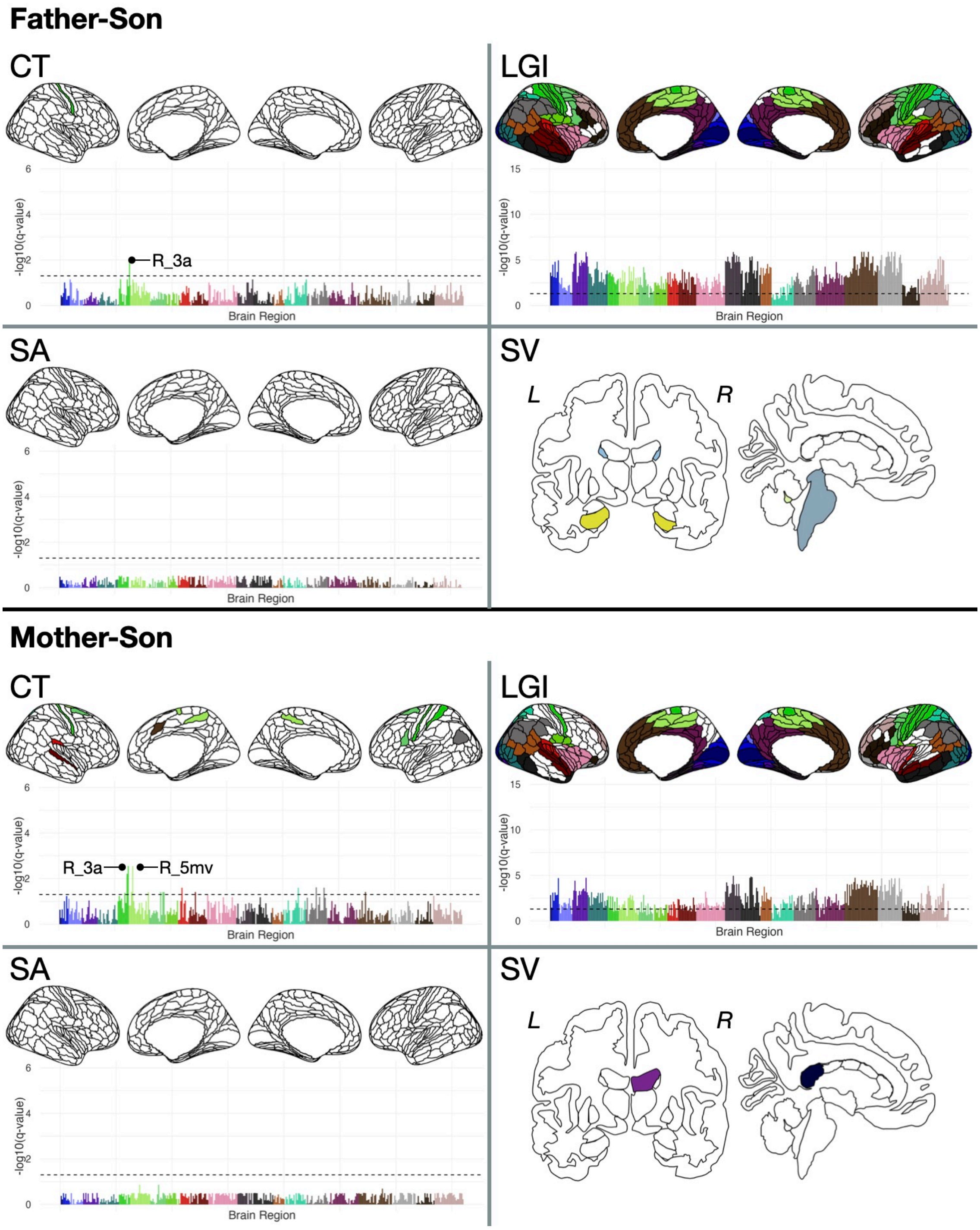


**Supplementary Figure 1. Neural similarities in father–son dyads and mother–son dyads**
The original HCP-MMP1 atlas (46) comprises 180 regions per hemisphere, grouped into 22 larger sections. The regions for which significantly stronger correlations were observed in father–son or mother–son dyads compared to unrelated pairs are color–coded. The *q*-values for the 360 brain regions are shown in the plot. The vertical axis of the plot represents the log-transformed *q*-values. Each bar represents an individual brain region, and the colors of the bars correspond to those in the brain illustration. Subcortical regions where significantly stronger correlations between parents and offspring compared to unrelated pairs were observed are color-coded according to the ggseg plotting tool (53). CT, cortical thickness; SA, surface area; LGI, local gyrification index; SV, subcortical volume, L, left hemisphere; R, right hemisphere.

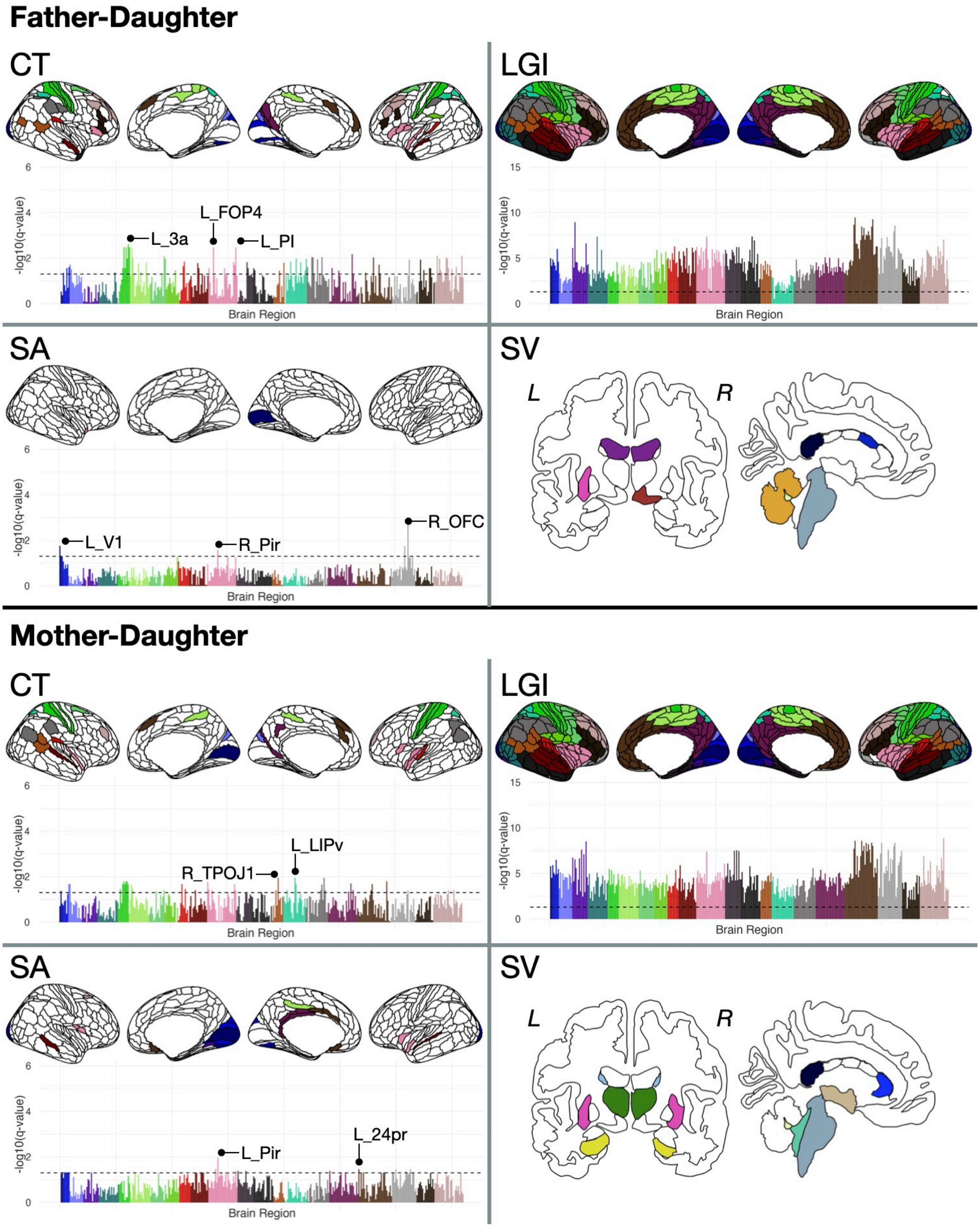


**Supplementary Figure 2. Neural similarities in father–daughter dyads and mother–daughter dyads**

The original HCP-MMP1 atlas (46) comprises 180 regions per hemisphere, grouped into 22 larger sections. The regions for which significantly stronger correlations were observed in father–daughter or mother–daughter dyads compared to unrelated pairs are color–coded. The *q*-values for the 360 brain regions are shown in the plot. The vertical axis of the plot represents the log-transformed *q*-values. Each bar represents an individual brain region, and the colors of the bars correspond to those in the brain illustration. Subcortical regions where significantly stronger correlations between parents and offspring compared to unrelated pairs were observed are color-coded according to the ggseg plotting tool (53). CT, cortical thickness; SA, surface area; LGI, local gyrification index; SV, subcortical volume, L, left hemisphere; R, right hemisphere.

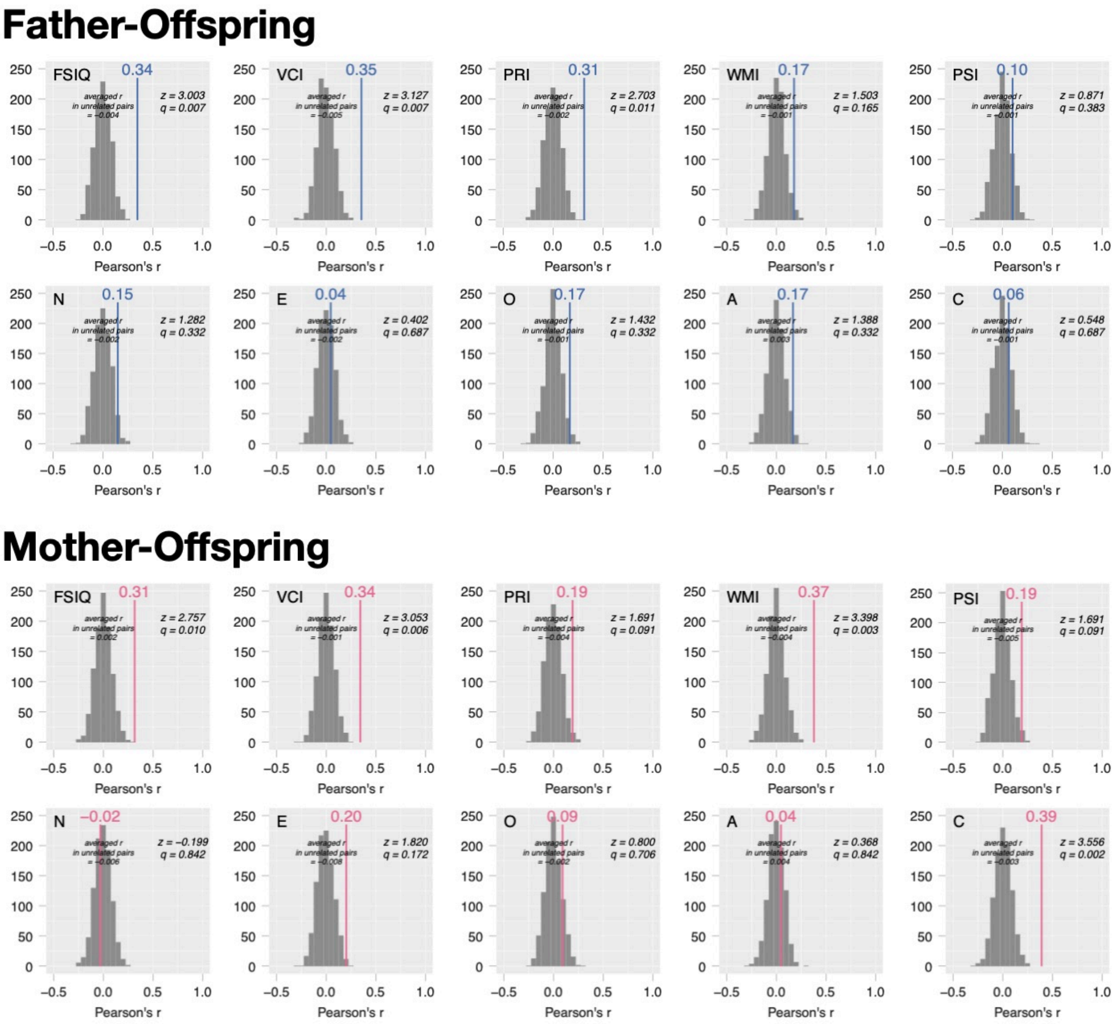


**Supplementary Figure 3. Parent–offspring behavioral similarities**

The differences in correlation coefficients between real parent-offspring dyads and unrelated pairs. The gray histogram represents the distribution of correlation coefficients for 1,000 patterns of unrelated pairs. “averaged r in unrelated pairs” means Z-transformed, averaged, and backtransformed correlation coefficients of unrelated pairs. The correlation coefficients for real parent–offspring pairs are shown as solid lines, with the corresponding values labeled on top. FSIQ, full-scale intelligence quotient; VCI, verbal comprehension index; WMI, working memory index; C, conscientiousness.

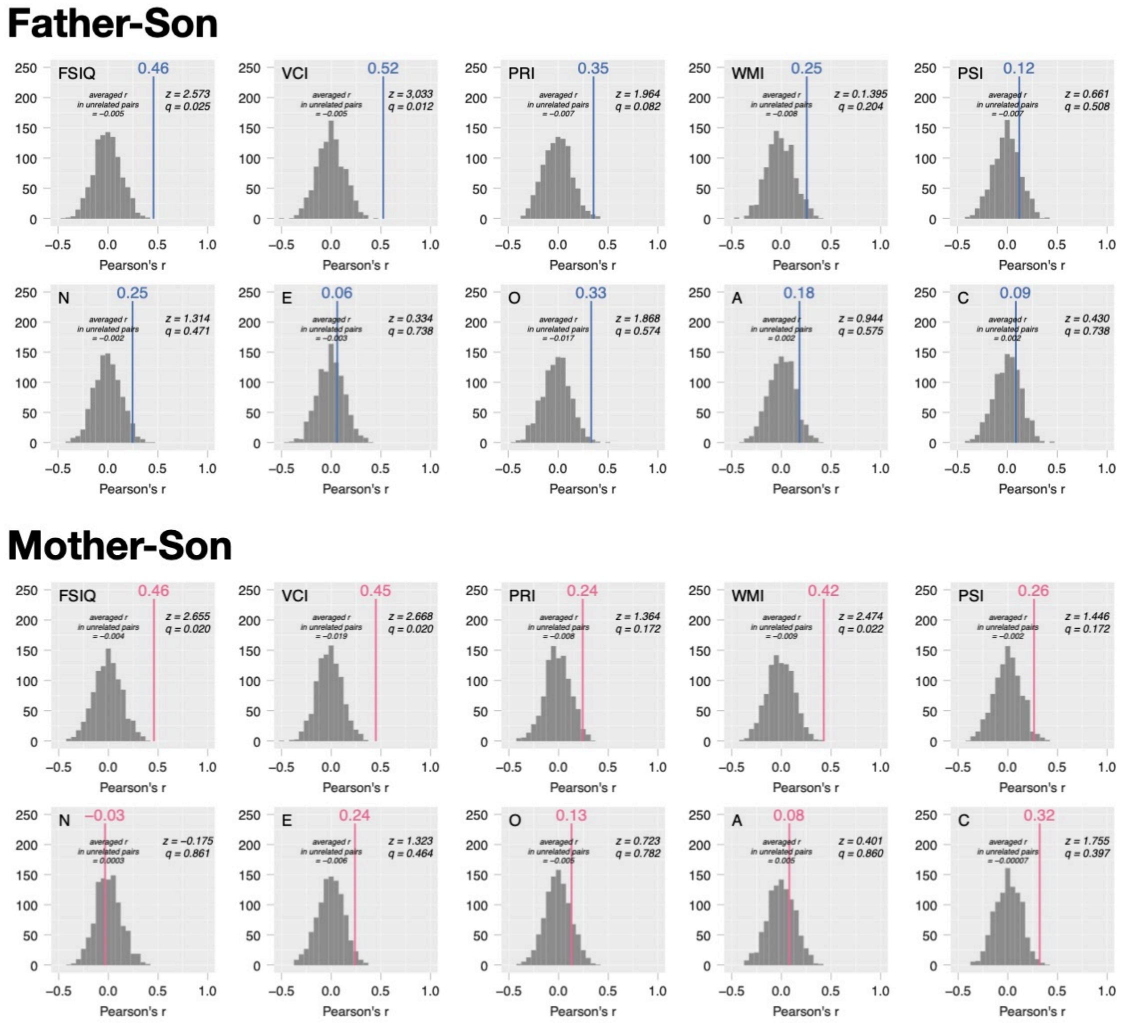


**Supplementary Figure 4. Behavioral similarities in father–son dyads and mother–son dyads**

The differences in correlation coefficients between real parent-son dyads and unrelated pairs. The gray histogram represents the distribution of correlation coefficients for 1,000 patterns of unrelated pairs. “averaged r in unrelated pairs” means Z-transformed, averaged, and backtransformed correlation coefficients of unrelated pairs. The correlation coefficients for real parent–offspring pairs are shown as solid lines, with the corresponding values labeled on top. FSIQ, full-scale intelligence quotient; VCI, verbal comprehension index; WMI, working memory index; C, conscientiousness.

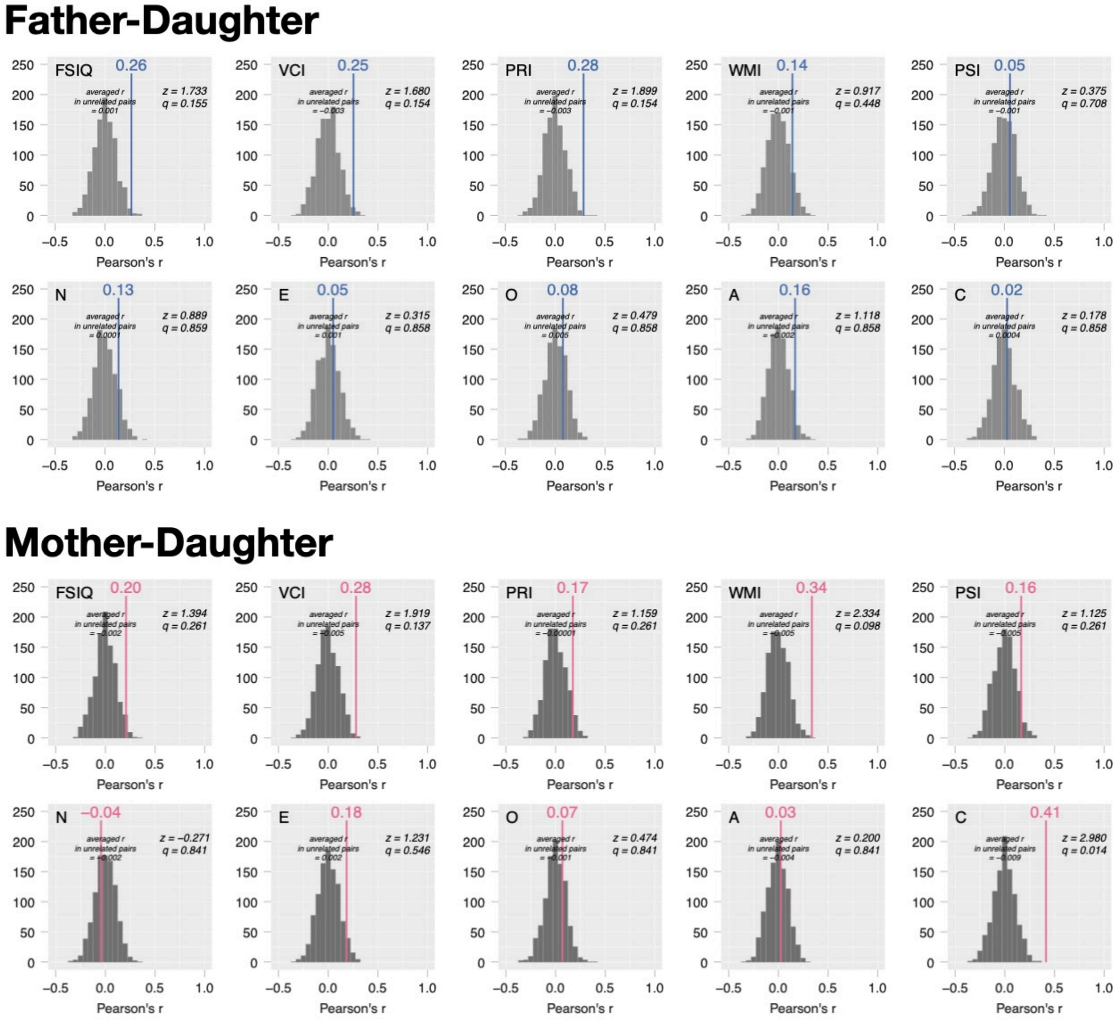


**Supplementary Figure 5. Behavioral similarities in father–daughter dyads and mother–daughter dyads**

The differences in correlation coefficients between real parent-daughter dyads and unrelated pairs. The gray histogram represents the distribution of correlation coefficients for 1,000 patterns of unrelated pairs. “averaged r in unrelated pairs” means Z-transformed, averaged, and backtransformed correlation coefficients of unrelated pairs. The correlation coefficients for real parent–offspring pairs are shown as solid lines, with the corresponding values labeled on top. FSIQ, full-scale intelligence quotient; VCI, verbal comprehension index; WMI, working memory index; C, conscientiousness.
